# *GNI-A1* mediates trade-off between grain number and grain weight in tetraploid wheat

**DOI:** 10.1101/540096

**Authors:** Guy Golan, Idan Ayalon, Aviad Perry, Gil Zimran, Toluwanimi Ade-Ajayi, Assaf Mosquna, Assaf Distelfeld, Zvi Peleg

**Affiliations:** The Robert H. Smith Institute of Plant Sciences and Genetics in Agriculture, The Hebrew University of Jerusalem, Rehovot 7610001, Israel; School of Plant Sciences and Food Security, Tel Aviv University, Tel Aviv 6997801, Israel

**Keywords:** Durum wheat, floret fertility, *Grain Number Increase 1*, grain weight, trade-off, wild emmer wheat

## Abstract

Grain yield is a highly polygenic trait determined by the number of grains per unit area, as well as by grain weight. In wheat, grain number and grain weight are usually negatively correlated. Yet, the genetic basis underlying trade-off between the two is mostly unknown. Here, we fine-mapped a grain weight QTL using wild emmer introgressions in a durum wheat background, and showed that grain weight is associated with the *GNI-A1* gene, a regulator of floret fertility. In-depth characterization of grain number and grain weight indicated that suppression of distal florets by the wild emmer *GNI-A1* allele increase weight of proximal grains in basal and central spikelets due to alteration in assimilate distribution. Re-sequencing of *GNI-A1* in tetraploid wheat demonstrated the rich allelic repertoire of the wild emmer gene pool, including a rare allele which was present in two gene copies and contained a non-synonymous mutation in the C-terminus of the protein. Using an F_2_ population generated from a cross between wild emmer accessions Zavitan, which carries the rare allele, and TTD140, we demonstrated that this unique polymorphism is associated with grain weight, independent of grain number. Moreover, we showed, for the first time, that *GNI-A1* proteins are transcriptional activators and that selection targeted compromised activity of the protein. Our finding expand the knowledge of the genetic basis underlying trade-off between key yield components and may contribute to breeding efforts for enhanced grain yield.

## Introduction

Wheat (*Triticum* sp.) is one of the major crops grown today, with production estimated at ~770 million tons per annum (http://www.fao.org/faostat). To meet the rising demand of the projected population growth by 2050, an increase of at least 60% in wheat production is required (Ray et al. 2013). Grain yield is a multifactorial trait determined by grain number (GN) and grain weight (GW), two primary yield components which are usually negatively correlated (Acreche and Slafer 2006; Bustos et al. 2013; García et al. 2013; Miralles and Slafer 1995; Sadras 2007; Slafer and Miralles 1993). GN is largely determined by the fertility of each floret within a spikelet (i.e. floret fertility). Following initiation of floret primordia, a large proportion of the florets will undergo degeneration through a genetically controlled, environment-responsive process of floret abortion (Ferrante et al. 2012; Ghiglione et al. 2008; Guo et al. 2017; Miralles et al. 1998; Prieto et al. 2018a; Prieto et al. 2018b; Sakuma et al. 2019), which defines the final number of grains per spikelet. From an evolutionary perspective, this process, which promotes high plasticity in floret/grain number, allows variable resource availability and a relatively stable phenotypical GW range (Sadras 2007; Sadras and Denison 2016). While GW is under the control of a complex genetic network expressed at various developmental stages, it is considered a very stable yield component with relatively high heritability. Final GW is largely affected by the duration and rate of linear grain growth and is a result of the interplay between potential grain weight (sink) and the actual supply of assimilates per grain during grain filling (source) (Fischer 2011).

Crop plants harbor only a small portion of the intra-species genetic diversity. The reduced genetic diversity in domesticated plants as compared to that of their progenitors (e.g. Harlan 1992; Haudry et al. 2007) is due to the limited number of founder genotypes during domestication (Eyre-Walker et al. 1998; Mayr 1942), coupled with subsequent selection for agronomic traits, referred to as evolution under domestication (Abbo et al. 2014; Ladizinsky 1998). Experimental data have shown that GW increased during wheat evolution from its direct progenitor, wild emmer [*T. turgidum* ssp. *dicoccoides* (Körn.) Thell.] (Abbo et al. 2014; Gegas et al. 2010; Golan et al. 2015). In parallel, an increase in floret fertility (i.e., number of grains per spikelet) was evident (Sakuma et al. 2019), that together with the increase in GW, promoted grain production in modern wheat. Recently, the quantitative trait locus *Grain Number Increase 1* (*GNI1*) was identified and characterized as a homeodomain leucine zipper class I (HD-Zip I) transcription factor. Reduced function mutation (N105Y) within the conserved homeodomain of *GNI-A1,* and knockdown of *GNI1* in transgenic hexaploid wheat, indicated that it is a suppressor of floret fertility. Transcript abundance of *GNI-B1* (orthologous copy on B genome) was negligible in floral organs of tetraploid and hexaploid wheat, and was suggested to be pseudogenized in the genome of ancestral *Aegilops* species (Sakuma et al. 2019).

Potential GW, defined as the intrinsic capacity of the grain to accumulate dry matter (Bremner and Rawson 1978), is relatively low in distal grain positions. Several studies indicated that the negative relationship between average GN and GW is an outcome of the high proportion of low potential GW in high yielding cultivars (Acreche and Slafer 2006; Ferrante et al. 2015; Fischer 2008; Miralles and Slafer 1995). Alternatively, Brenner and Rawson (Bremner and Rawson 1978) showed that distal grains have similar potential GW as proximal grains. In addition, removal of florets prior to grain filling increased weight of the remaining grains (Calderini and Reynolds 2000b; Fischer and HilleRisLambers 1978), suggesting a degree of growth limitation imposed by competition for insufficient source.

The physiological mechanisms underlying the trade-off between GN and GW have been intensively studied, however, little is known about the genetic basis underlying this trade-off in wheat. Previously, we mapped a major QTL affecting GW on chromosome 2AL, with the wild emmer allele conferring heavier grains (Golan et al. 2015). Here, we re-examined the effect of this QTL on GW and its association with additional yield components. Subsequently, we fine-mapped the GW QTL and examined its genetic association with the *GNI-A1* gene. Detailed characterization of the phenotype was conducted to elucidate the physiological mechanisms linking the two loci in a narrow genomic region. Moreover, we used wild and domesticated tetraploid genepools to explore the allelic variation in the *GNI-A1* gene and investigate its effect on gene function. Our findings shed new light on the interaction between yield components during wheat evolution under domestication.

## Materials and methods

### Growth conditions

Characterization of yield components, QTL mapping and fine-mapping were carried out in an insect-proof screen-house (0.27 × 0.78 mm pore size screen) at the experimental farm of the Hebrew University of Jerusalem in Rehovot, Israel (34°47′ N, 31°54′ E; 54 m above sea level). The soil at this location is brown-red, degrading sandy loam (Rhodoxeralf) composed of 76% sand, 8% silt and 16% clay. Plants were treated with pesticides to protect against pathogens and insect pests and the plots were weeded, by hand, once a week.

Characterization of GW in the F_2_ (Zavitan × TTD140) population was conducted in Atlit experimental station (32°41′14″N 34°56′18″E; 22m above sea level). The soil at this location is a brown soil, composed of 18.7% sand, 22.6% silt, 57.0% clay and 1.7% organic matter.

### Phenotypic measurements

The number of spikelets per spike (StPS), grains per spike (GPS) and GW, were obtained from a sample of three spikes for QTL mapping and five spikes for yield components characterization and fine-mapping of GPSt and GW. GPSt was derived by dividing GPS by StPS. GW derived by dividing the total GW of each spike by GPS. Grain weight per plant (GWPP) derived from the total GW of each plot divided by the number of plants.

### QTL mapping

Re-examination of the GW QTL and QTL mapping of grain number per spike (GPS) and grain number per spikelet (GPSt) was conducted using a RISL (Recombinant Inbred Substitution Lines) population derived from a cross between durum cultivar Langdon (LDN) and the chromosome substitution line ‘DIC-2A’, which harbors the 2A chromosome of wild emmer accession FA-15-3 (ISR-A) in the homozygous background of LDN (Cantrell and Joppa 1991; Joppa 1993). For QTL mapping, we applied a randomized complete block design with three replicates (single plants). Linkage analyses and map construction were performed based on 30 genetic markers located on chromosome 2A (Table S1), using the evolutionary strategy algorithm included in the MultiPoint package, as previously described (Golan et al. 2015). QTL analysis was performed with the MultiQTL package (http://www.MultiQTL.com), using the general interval mapping for Recombinant Inbred Lines (RIL populations. The hypothesis that one locus on the chromosome has an effect on a given trait (H_1_) was compared with the null hypothesis (H_0_) that the locus has no effect on that trait. Once the genetic model was chosen, 10,000 bootstrap samples were run to estimate the standard deviation of the main parameters: locus effect, its chromosomal position, its LOD score and the proportion of explained variation (PEV).

### Fine mapping

For fine-mapping, three RISLs (#4, #63, #102), each of which harbored the DIC-2A allele of a marker defining the *GW* QTL (Golan et al. 2015), were backcrossed to LDN and then self-pollinated to produce three independent F_2_ populations. To identify recombinants within the QTL interval, 1006 F_2_ progenies were genotyped with microsatellite markers *Xgwm558* and *Xcfa2043,* located outside and in vicinity to the proximal and distal ends of the QTL. In the following generation, eight F_3_ individuals per F_2_ plant were genotyped with *Xgwm558*, *Xcfa2043* and *Xhbg494* which was the closest marker to QTL peak. Recombinant lines were further genotyped with 12 additional markers (Table S1) to identify homozygous recombinants within the QTL region. Eighteen F_4_ backcrossed recombinant lines were used for fine-mapping of GPSt and GW. A split plot, complete random block design (*n*=5) was applied; each plot (1×1 meter) contained twenty plants per line.

### Phenotypic and genetic characterization of F2 progenies

Characterization of GW in F_2_ was conducted using 252 progenies derived from a cross between wild emmer accessions Zavitan and TTD140. Following anthesis, spikes were covered with glassine grain bags to avoid grain dispersal. A single spike per plant was evaluated for GN and GW. Allele determination of *GNI-A1* locus was conducted using the *Xhuj11* marker (Table S1).

### Physiological characterization

Detailed characterization of the *GNI-A1* effect was conducted using LDN and a backcrossed recombinant line (63-18) which carries a wild emmer *GNI-A1* allele in a relatively small introgression (~75Mbp) and holds recombination within the QTL interval. Spikelet removal or distal floret removal was imposed at heading in basal (lower third of the spike) or central (middle third of the spike) spikelets (see Fig. 1) Four spikes per plot (*n*=5) were harvested at maturity and spikelets were manually dissected to obtain number of grains per spikelet and their corresponding GW. The effect of spikelet and floret removal was tested using comparisons between measurement of GPSt and GW in treated *vs*. untreated plants.

To examine the effect of *GNI-A1* on grain development, grains were sampled at heading, anthesis, and 8, 14, 21, 28 and 39 days after anthesis. Four basal, central and apical spikelets were dissected from synchronously flowering spikes (*n*=3). Grains were oven dried at 60°C for 5 days before weighing.

### Haplotype analysis

A haplotype analysis was conducted based on re-sequencing data of *GNI-A1* (5'←3', 600bp) from 47 wild emmer, 28 domesticated emmer and 36 durum cultivars (Table S2). The re-sequencing data combines 72 accessions reported in Sakuma et al. (2019) with 39 additional accessions. Multiple sequence alignments were performed using ClustalOmega (Sievers and Higgins 2014). Phylogenic tree construction was conducted using the *phylogeny.fr* web tool (Dereeper et al. 2008).

### Copy number estimation

Genomic DNA was extracted from fresh leaf tissue of individual seedlings using the C TAB protocol. qPCR was carried out, using PerfeCTa SYBR Green FastMix (Quanta Biosciences Inc.), on the PikoReal RT-PCR system (Thermo Fisher Scientific Inc.). An efficiency value of 100±10% was confirmed for both primer sets (Table S1). DNA samples were diluted to 25ng/μL, based on a standard curve of five serial dilution points. Samples were denatured at 95° C for 3 min, followed by 40 cycles at 95° C for 10 sec and 60° C for 45 sec. The 2^−ΔΔCT^ method (Livak and Schmittgen 2001) was used to normalize and calibrate copy number of *GNI-A1* relative to the single copy gene *TraesCS2A02G134000*.

### Transcriptional activity in yeast

Coding sequences of Hap6 and Hap9 were PCR-amplified from cDNA using primers extended with EcoRI and SalI recognition sequences. Resulting PCR products were cloned into EcoRI/SalI-digested pBD-GAL4 (Clontech, CA, USA) using standard restriction ligation. Following ligation, plasmids were transformed into *Escherichia coli* (DH5a) using the heat shock method and then sequence-verified. Site-directed mutagenesis was carried out using the QuikChange Lightning Multi Site-Directed Mutagenesis Kit (Agilent, CA, USA) according to manufacturer's instructions.

The 5'- GCGTGTGCGGCGGGAGCCC-GGGCTCATCCTTCTCGA-3' mutagenic primer was used to introduce N180S into pBD-Hap6 (resulting in pBD-Hap3). 5'-CCGTTAGCGTGTGCGGCGGGAGCCCGCGCTCATCCTTCTCGACGGGA-3' was used similarly to introduce both N180S and G182R (resulting in pBD-Hap7). Finally, The 5'-TCCCGCCGCACACGCTAACGGCGCCAGCGGAGTCTCCGCCATCGGCGTGGATCCCAGCTGCCG-3' primer was used to introduce N166D to the pBD-Hap3 (pBD-Hap8).

All pBD plasmids were individually transformed into the Y190 yeast strain using the LiAc/SS Carrier DNA/PEG Method (Xiao 2006). pBD-X harboring clones were isolated by growth on synthetic dextrose (SD) agar plates lacking the amino acid Tryptophan. Twenty transformed clones from each line were pooled and 3μl drops were spotted on new SD–Trp plates. Following an additional growth phase, a β-galactosidase activity assay was carried out to monitor *LacZ* expression levels. Yeast were chloroform-lysed and stained, as previously described (Park et al. 2009).

Prediction of phosphorylation sites on GNI-A1 was conducted using NetPhos 3.1 (Blom et al. 1999; Blom et al. 2004).

### Statistical analysis

All statistical analyses were conducted using the JMP^®^ ver. 14 statistical package (SAS Institute, Cary, NC, USA). Principal component analysis (PCA) was used to determine the associations between yield components. PCA was based on a correlation matrix and is presented as biplot ordinations of the backcrossed recombinant lines (PC scores). Two components were extracted using eigenvalues >1.5 to ensure meaningful implementation of the data by each factor. Correlations between yield components in backcrossed recombinant lines were estimated using the Row-wise method, using mean values for each genotype. To assess the allelic effect on GPSt and GW, mean values of experimental units were analyzed using one-way ANOVA, assuming a two-tailed distribution and equal variance. Components of descriptive statistics are graphically presented in box plots: median value (horizontal short line), quartile range (rectangle) and data range (vertical long line).

## Results

### *GNI-A1* locus regulates grain number and grain weight

To examine how the GW locus affects grain production, grain yield components were characterized in two field experiments (14 RISL in 2016 and 18 backcrossed recombinant lines in 2017). PCA extracted two major components, explaining collectively 77% of the phenotypic variance among the backcrossed recombinant lines. Principal component 1 (PC1; X axis) explained 50.9% of the dataset variance and was loaded positively with the number of spikes per plant (SPP), spikelets per spike (StPS), grains per spike (GPS), grains per spikelet (GPSt) grain weight per plant (GWPP) and negatively loaded with GW. PC2 (Y axis) explained 26% of the variance and was positively loaded with SPP, GWPP and GW and negatively loaded with GPS, GPSt and StPS (Fig. 1a). GW was negatively correlated with GPSt and GPS (*P*<10^−4^) with correlation coefficients being r=0.7 and 0.6, respectively (Table S3).

**Fig. 1.**
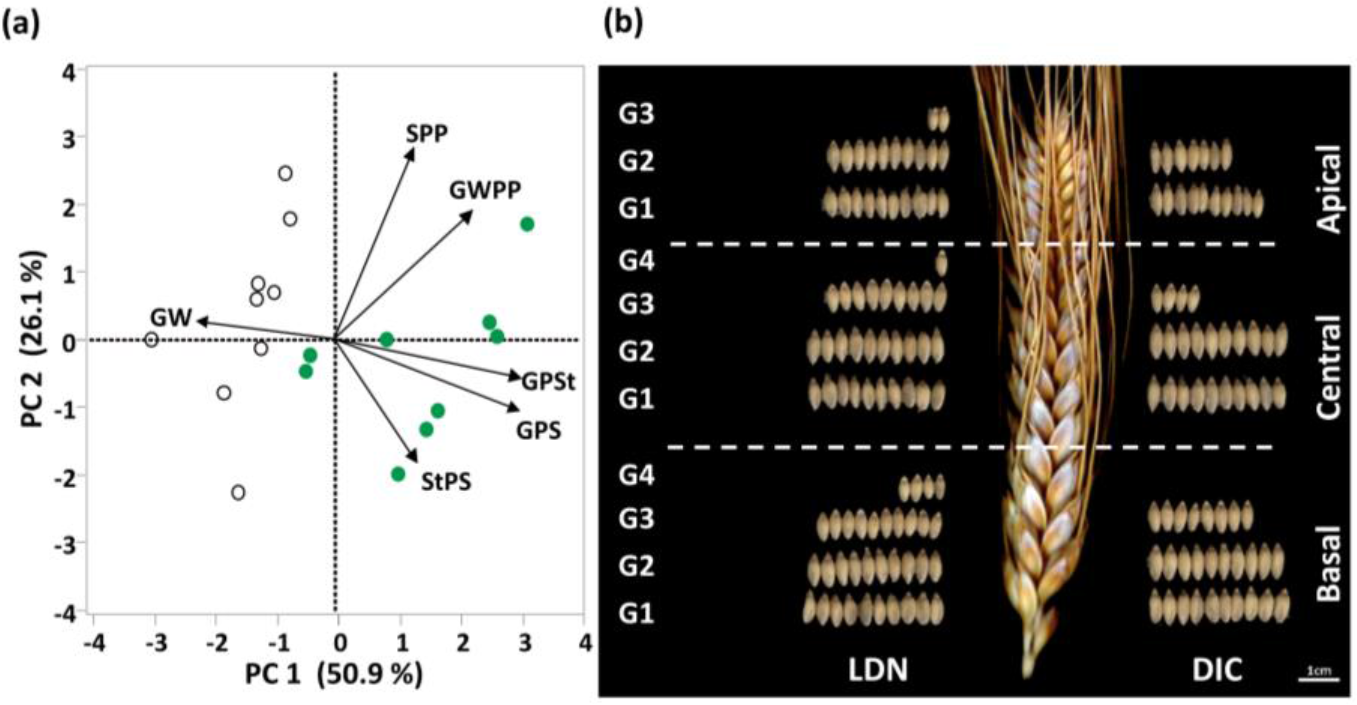
The effect of grain weight QTL on grain yield components. **a)** Principal component analysis of continuous plant traits in 18 backcrossed recombinant lines. Biplot vectors are trait factor loadings for principal component (PC) 1 and PC2. Green and white dots represent genotypes carrying the domesticated (LDN) and wild (DIC) allele, respectively. Grain weight (GW), spikes per plant (SPP), grain weight per plant (GWPP), grains per spikelet (GPSt), grains per spike (GPS) and spikelets per spike (StPS). **b)** Demonstration of the locus effect on floret fertility and grain size in representative basal, central and apical spikelets.

The wild emmer (DIC) allele defined by the *Xhbg494* marker (Golan et al. 2015) was associated with higher GW, while the durum (LDN) allele was associated with high GPSt and GPS (Fig. 1b, Supplementary Fig. S1). The number of spikelets per spike was not affected by the GW locus and therefore higher GPS in LDN derived directly from GPSt (r=0.96, *P*<0.0001). Genotypes carrying either LDN or DIC alleles had similar plant height (177 cm and 180 cm for DIC and LDN, respectively; *P*=0.5) and days to flowering (on average 133 days). The increase in GW by the DIC allele compensated for the lower GPS and GPSt and maintained similar grain yield between alleles in the backcrossed recombinant lines. Notably, in RISL, GWPP was slightly higher in LDN genotypes (*P*=0.06) and was associated with SPP (Supplementary Fig. S1, Table S3).

Genetic analysis using a RISL population showed a stable chromosomal position of the GW QTL in two field trials, in 2014 (Golan et al. 2015) and 2017 (this study), which explained 48% and 63% of the phenotypic variation, respectively (Supplementary Table S4). Subsequently, QTL mapping of GPS and GPSt showed these traits co-localize with the GW QTL and explained 54% and 61% of the phenotypic variance, respectively, (Fig. 2a, Table S4). To fine-map the QTL and dissect the genetic relationship between GPSt and GW, we used backcrossed recombinant lines and mapped the QTLs as a simple Mendelian locus. GW and GPSt (*GNI-A1*) loci did not recombine and were delimited within a 5.4 Mbp interval (Fig. 2b), which is associated with a significant trade-off between these yield components (Fig. 2b, c). The *GNI-A1* locus contains the *GNI-A1* gene. Genotypes carrying the DIC allele harbor a functional *GNI-A1*, whereas in LDN, amino acid substitution N105Y in the conserved homeodomain, impairs gene function and increases floret fertility (Sakuma et al. 2019).

**Fig. 2.**
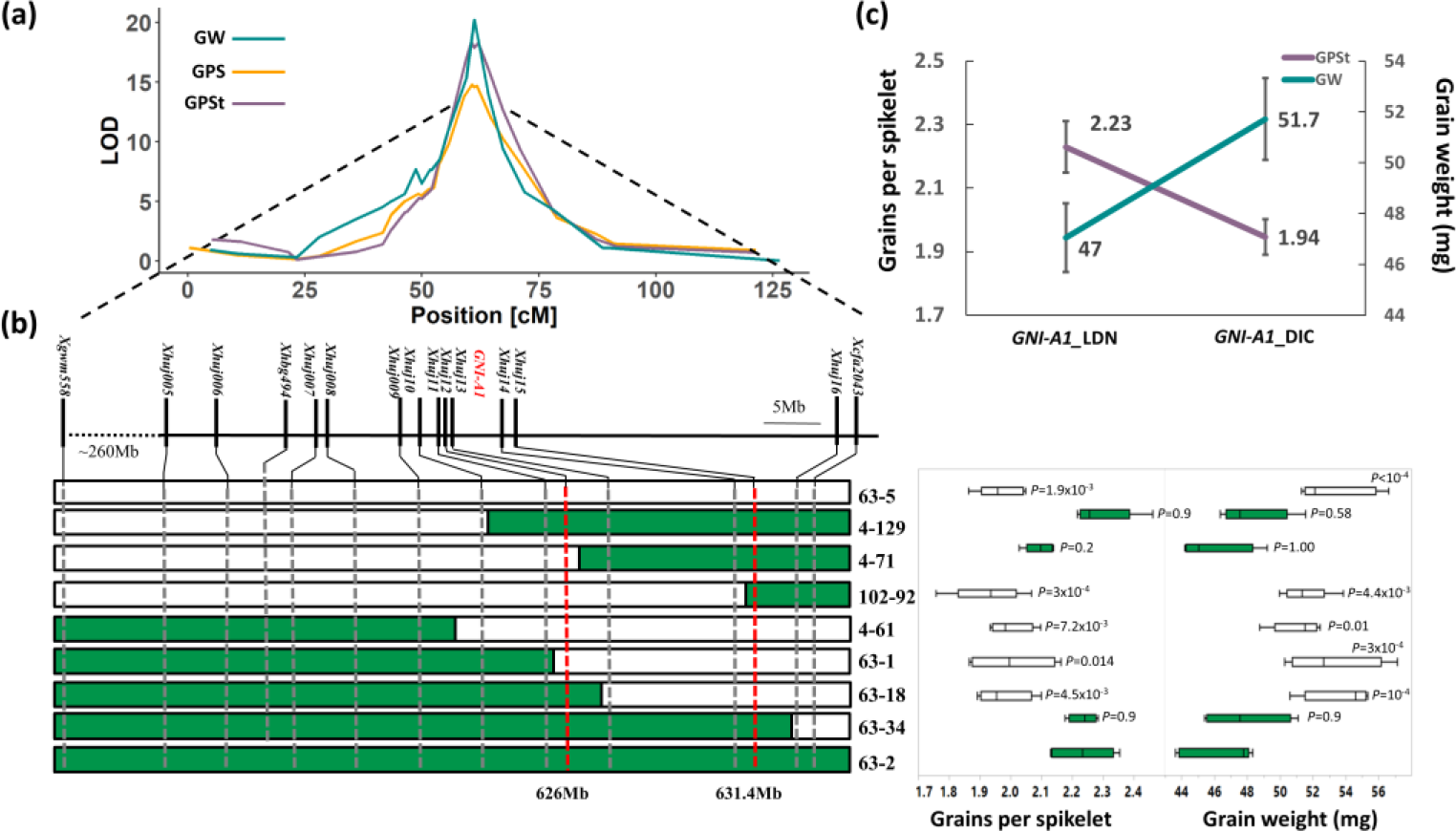
Fine-mapping of grain number and weight. **a**) QTL mapping of grains per spike (GPS), grains per spikelet (GPSt) and grain weight (GW) on chromosome 2A. **b**) Graphical genotyping demonstrating fine-mapping of GW and GPSt using backcrossed recombinant lines. *P* values indicate differences from 63-2, a backcrossed recombinant line carrying domesticated allele, along the mapping interval, as determined by Dunnett’s test. Red dashed lines indicate GPSt (*GNI-A1*) and GW loci intervals. **c**) Significant trade-off between GPSt and GW in backcrossed recombinant lines carrying domesticated (*GNI-A1_*LDN) and wild emmer (*GNI-A1_*DIC) alleles (*n*=9).

### Reduction in grain number is spatially associated with enlarged grains

To study the spatial effect of the *GNI-A1* locus, GN and the weight of grains across all spikelet positions were measured in LDN and a backcrossed recombinant line (#63-18) which carries the *GNI-A1*_DIC allele in a LDN background. Basal spikelets had more grains in LDN and DIC genotypes as compared to central and apical spikelets, which carried the lowest number of grains per spikelet (Fig. S2a). Introgression of the DIC allele significantly decreased the GN in basal (2.8 *vs*. 3.3; *P*<0.0001) and central (2.0 *vs*. 2.6, *P*<0.0001) spikelets, with a non-significant effect in apical spikelets (Fig. 3a, Supplementary Fig. S2a).

The spatial effect of *GNI-A1* on GW was related to the number of grains per spikelet (Fig. 3, Supplementary Fig. S2). The DIC allele increased GW of G1 (indicates grains produced at the 1^st^ floret) and G2 (indicates grains produced at the 2^nd^ floret) in basal and central spikelets, but had no significant effect on GW in apical spikelets and distal florets (i.e., G3) (Fig. 3b, Supplementary Fig. S2b). Characterization of grain development indicated that the effect of *GNI-A1* on GW manifested during late stages of grain filling (Supplementary Fig. S3). Moreover, the suppression of distal florets by the DIC allele decreased their proportion in the final GN (Supplementary Fig. S4), thus, contributing to the increase in average GW.

**Fig. 3.**
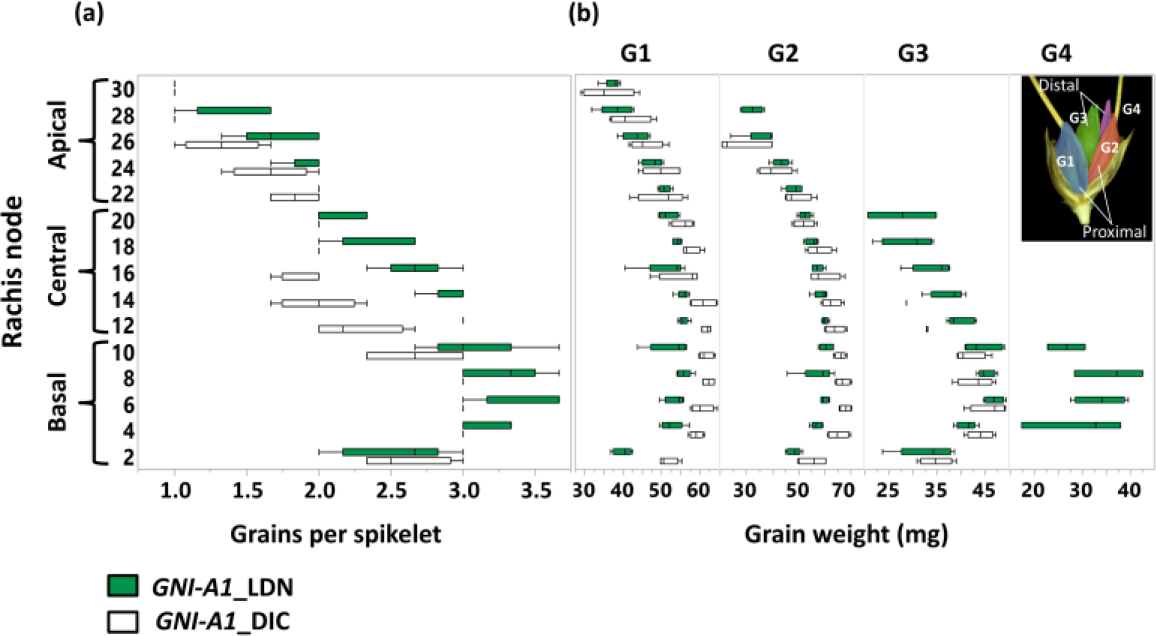
Phenotypical characterization of the *GNI-A1* effect on grains per spikelet (GPSt) and grain weight (GW) in main shoots of durum cultivar Langdon (LDN) and a backcrossed recombinant line (63-18) containing the wild (*GNI-A1*_DIC) allele in the background of LDN. **a**) Number of grains per spikelet and **b**) weight of particular grain positions along the spike. Values are means of four spikes per plot (*n*=5). *GNI-A1_LDN* and *GNI-A1*_DIC alleles are represented by green and white box plots, respectively.

### Competition among sinks is likely to reduce grain weight in LDN background

To examine if the trade-off between GN and GW in basal and central spikelets is due to competition among grains for insufficient source, we manipulated GN by removal of either basal or central spiklets, thereby altering assimilates distribution within the spike. In LDN, either basal or central spikelet removal increased weight of G1, G2 and G3 in basal, central and apical spikelets (Fig.4a, Supplementary Fig. S5a).

**Fig. 4.**
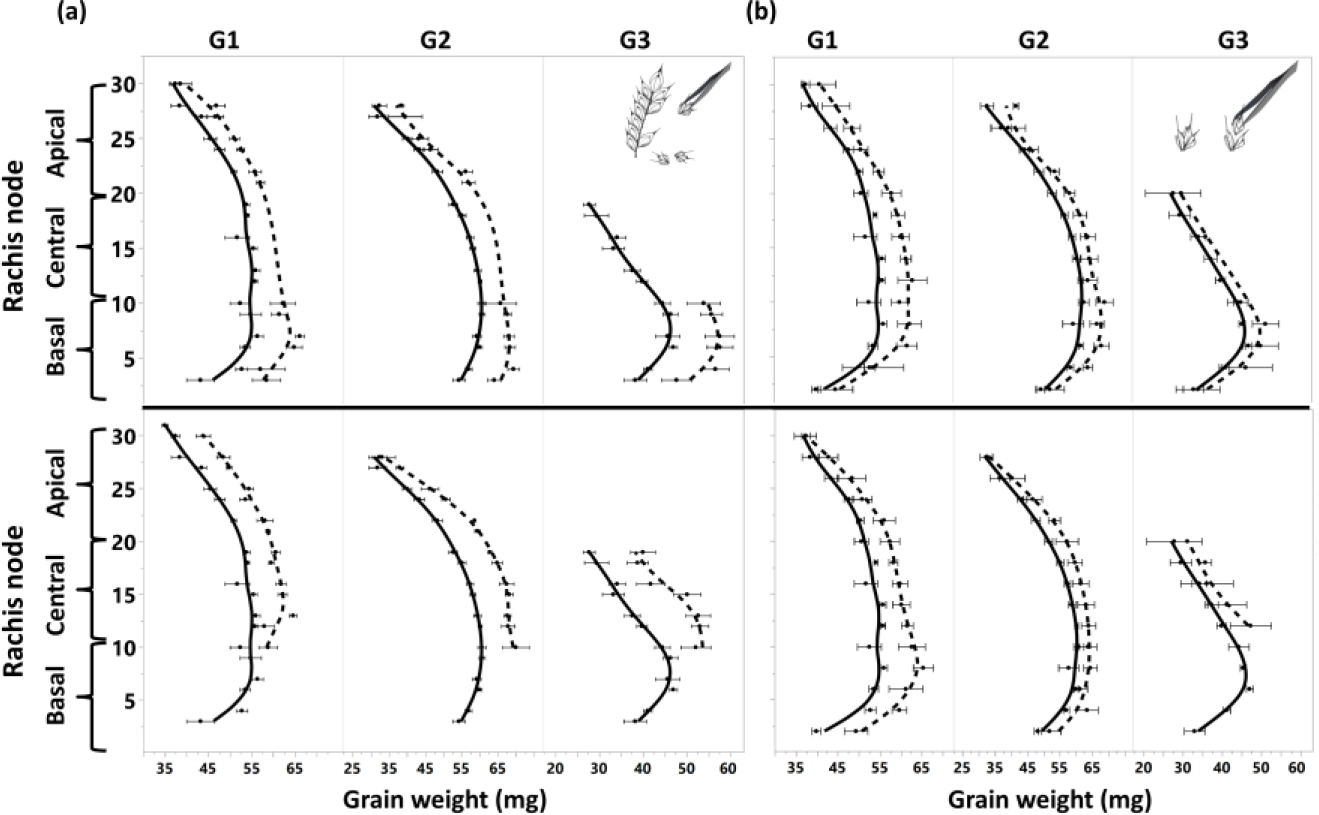
Effect of source/sink modifications and spikelet position on weight of particular grain positions along the LDN spike. **a)** GW following removal of basal (lower) or central (upper) spikelets. **b)** GW following removal of distal florets in basal (lower) or central (upper) spikelets. Solid and dashed lines represent control and treated spikes, respectively. Values are mean±SE (*n*=5).

In the backcrossed recombinant line, the effect of spikelet removal was weaker and only seldom expressed (Supplementary Fig. S5), indicating that LDN is more source-limited as compared. In order to imitate the GN phenotype in plants carrying the DIC allele, we removed only the distal florets in basal or central spikelets of LDN. Generally, removal of distal florets increased weight of G1 and G2 (Fig. 4b, Supplementary Fig.S6), indicating that a high proportion of distal grains imposes source limitation for the growth of G1 and G2. Consistent with spikelet removal, the effect of floret removal on GW was greater in LDN. Removal of distal florets in basal spikelets increased weight of G1 (6.37 mg, *P*=0.03 *vs*. 5.00 mg, *P*=0.03) and G2 (5.25 mg, *P*=0.02 *vs*. 4.00 mg, *P*=0.4) in LDN *vs*. DIC, respectively. Removal of florets in central spikelets increased weight of G1 and G2 in LDN (5.87 mg, *P*=0.07 and 5.00 mg, *P*=0.04, respectively) but had no significant effect on GW in the backcrossed recombinant line. The effect of floret removal was not restricted to treated spikelets and extended to remote spikelets, suggesting that presence of G3 was not a physical restraint on growth of G1 and G2 (Supplementary Fig.S6). Overall, our findings suggest that lower spikelet fertility associated with the *GNI-A1*_DIC allele, eases competition among developing grains and increases proportion of high-weight grains. Therefore, we suggest that the *GNI-A1* gene is the likely candidate for the GW QTL.

### Allelic variation of *GNI-A1*

To study the origin and distribution of *GNI-A1* gene variants during wheat evolution, we re-sequenced the alleles present in 111 accessions of wild emmer, domesticated emmer and durum wheat. We identified eight haplotypes of wild emmer and two haplotypes of domesticated emmer and durum wheat, each. Four amino acid substitutions were found among haplotypes, including the N105Y reduced-function mutation within the conserved homeodomain, that exist only in durum wheat and was shown to increase floret fertility (Sakuma et al. 2019). Three non-synonymous amino acid substitutions (i.e., N166D, S180N and G182R) were found within the C-terminal region (CTR) (Fig. 5).

**Fig. 5.**
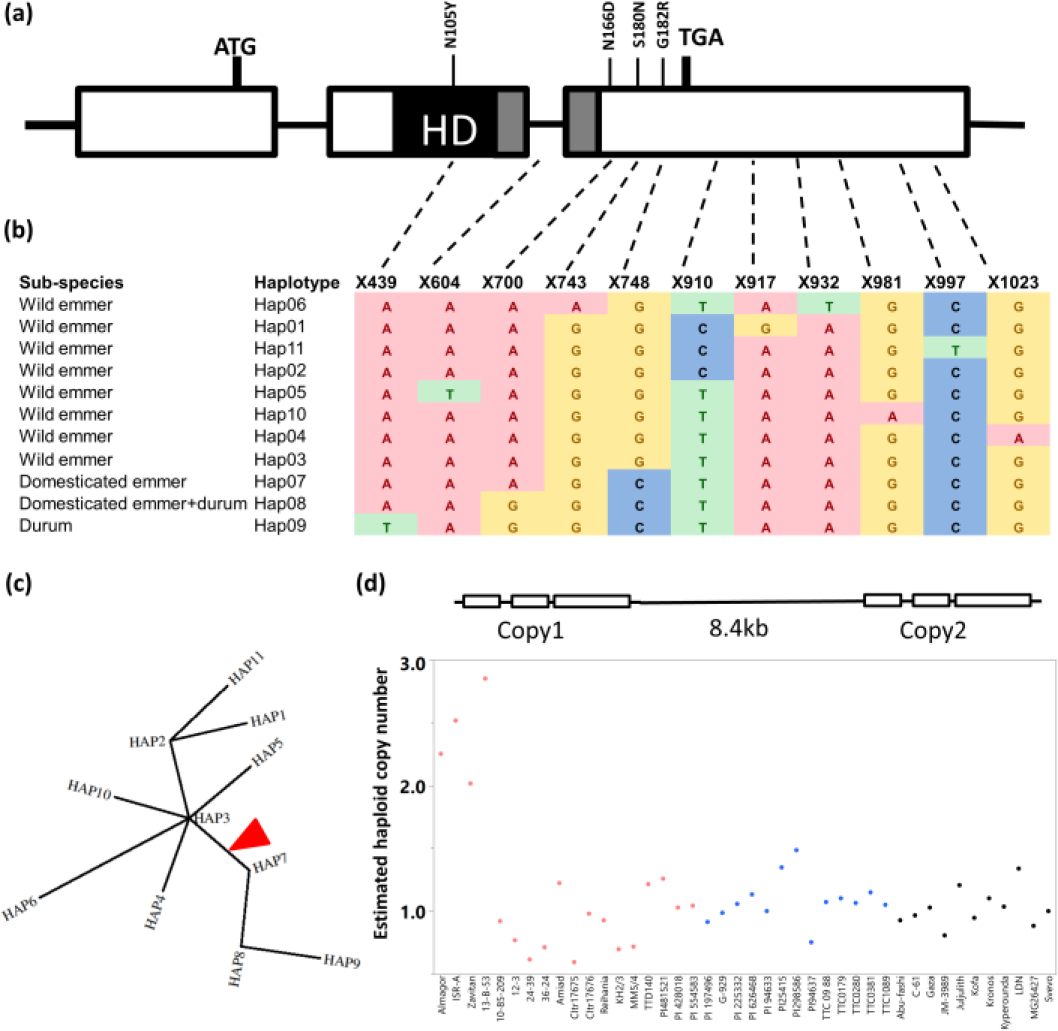
Haplotype analysis of *GNI-A1* in tetraploid wheat. **a**) Gene structure of *GNI-A1*. Exons are indicated as boxes. Variation in amino acid sequence is indicated above. Single Nucleotide Polymorphism (SNP) are connected to the haplotype table by dashed lines. X indicates nucleotide position from the start of the ORF. **b)** Eleven haplotypes classified by SNP variations were detected in tetraploid wheat. **c**) Phylogenetic analysis of the eleven haplotypes. Red triangle indicates wheat domestication on the phylogenetic tree. **d**) *GNI-A1* locus on chromosome 2AL of the wild emmer genome (accession Zavitan). Mean (*n*=3) qPCR estimates of *GNI-A1* haploid copy number based on *GNI-A1* quantity relative to *TraesCS2A02G134000*. Wild emmer wheat (Pink), domesticated emmer (blue) and durum wheat (black).

Haplotype analysis revealed that the S180N substitution is exceptional in wheat, confined to four wild emmer accessions from Northern Israel and absent from domesticated wheat genepool. The additional amino acid substitutions N105Y, N166D and G182R were evident only in domesticated wheat. G182R was introduced during the transition from Hap3 (wild emmer) to Hap7 (domesticated emmer) and was associated with the domestication of tetraploid wheat. N166D was introduced post-domestication (Hap8) and is present in domesticated emmer as well as durum cultivars (Fig. 5, Supplementary Table S2).

Investigation of *GNI-A1* in the wild emmer wheat genome (accession Zavitan; Avni et al. 2017) identified two copies of *GNI-A1* on the long arm of chromosome 2A (Fig. 5d). However, in the genomes of domesticated hexaploid wheat (*cv*. Chinese Spring; https://www.wheatgenome.org) and tetraploid durum wheat (*cv*. Svevo; Maccaferri et al. 2019), we detected only a single copy of *GNI-A1*. To further characterize variation in copy number among tetraploid genepool, we estimated copy number of *GNI-A1* in 44 accessions of wild and domesticated wheat using qPCR. A single copy of *GNI-A1* was estimated in all wild and domesticated accessions tested, with the exception of four wild emmer accessions from Northern Israel, corresponding to Hap6 (accessions: Zavitan, ISR-A, 13-B-53 and Alm-1; Supplementary Table S2), which were estimated to carry two or more copies (Fig. 5d).

### Allelic variation analysis reveals new sources for functional diversity of *GNI-A1*

In two-rowed barley, amino acid substitution (S184G) within the CTR of *VRS1* (*GNI-A1* barley homolog) alters phosphorylation potential of VRS1 and increases GW (Sakuma et al. 2017). Allelic variation analysis exposed a non-synonymous substitution (S180N) in a similar domain of GNI-A1, confined to wild emmer accessions (Hap6). (Figs. 5, 6a, Supplementary Table S2). To test whether the S180N amino acid substitution drives an increase in GW, we examined the association between the S180, N180 alleles and GW of 124 F_2_ genotypes homozygous for *GNI-A1*. The F_2_ population was developed from a cross between two wild emmer accessions, Zavitan (Hap6) and TTD140 (Hap4), both carrying a functional homeodomain (105N). Zavitan carries asparagine (N) in position 180 of GNI-A1 instead of canonical serine (S) found in TTD140 (Fig. 6a). Zavitan had more spikelets per spike and a higher number of grains per spike. GW did not differ significantly between parental lines (Fig. S7). Genetic analysis of spikes from F_2_ showed that the Zavitan allele is associated with higher GW (39 *vs*. 36 mg), regardless of grain number per spike (Fig. 6b,c). Prediction of phosphorylation sites on GNI-A1 showed that the S180N substitution resulted in hypo-phosphorylation (Supplementary Fig. S8) of the CTR.

**Fig. 6.**
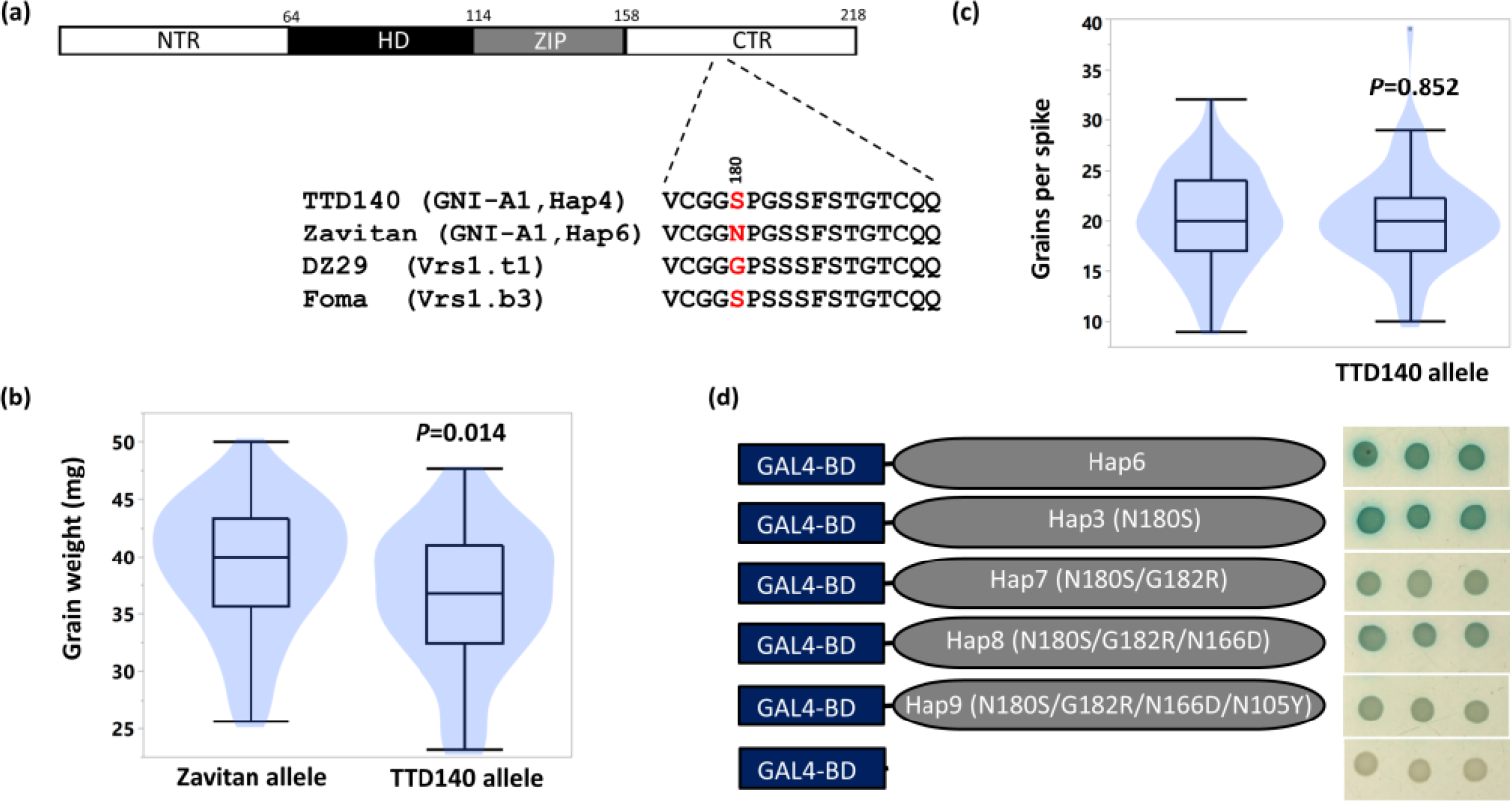
Functional analysis of amino acid substitutions in *GNI-A1*. **a)** Wild emmer accession Zavitan and DZ29 (two-rowed barley) encode an amino acid substitution from canonical serine (S) present in wild emmer TTD140 and barley *cv*. Foma, respectively. **b**) Association between the *GNI-A1* allele and grain weight and **c**) grain number per spike, in homozygous F2 lines derived from a cross between Zavitan and TTD140 (*n*=62). *P* values were determined using Student’s t-test. **c**) Transcriptional activity of *GNI-A1* in yeast. The *LacZ* gene reports transcriptional activity of the Zavitan allele (Hap6), mutated Hap6 alleles, where 180N was substituted to 180S (Hap3), 182G to 182R (Hap7), 166N to 166D and a domesticated durum allele (Hap9).

Reasoning that GNI-A1 might retain its activity in a heterologous system, a transcription activation assay was used to compare GNI-A1 alleles. The yeast GAL4 DNA-binding domain (GAL4BD) was fused to the open reading frame (ORF) of the wild emmer 180N allele (i.e., Zavitan, Hap6) and to mutated versions of this ORF; site-directed mutagenesis was conducted to substitute 180N to 180S, 182G to 182R and 166N to 166D. Furthermore, GAL4BD was fused to the ORF of durum cultivar LDN (Hap9). In the Y190 yeast strain, the *LacZ* reporter-gene is placed under a GAL1 upstream activating sequence (UAS), which is recognized by BD. While the unfused BD construct (negative control) didn’t induce reporter gene expression, transcriptional activity of GNI-A1 was evident in domesticated as well as in wild emmer proteins, implying that GNI-A1 is a transcriptional activator. Amino acid substitution at position 180 did not bear a substantial effect on transcriptional activity, suggesting that, at least in yeast, this substitution does not impact transcriptional activity of GNI-A1. Nevertheless, wild emmer alleles conferred higher transcriptional activity as compared to the domesticated emmer (Hap7, Hap8) and durum alleles (Hap8, Hap9) (Fig. 6d), suggesting that amino acid substitution G182R, associated with wheat domestication, led to decreased transcriptional activity of GNI-A1.

## Discussion

Wheat grain yield is a highly polygenic trait, with low heritability, influenced by various genetic, environmental and management factors (i.e., G×E×M) and is the outcome of the interaction between grain number per m^2^ and grain weight. Although the physiological mechanisms underlying trade-off between GN and GW have been extensively studied, the genetic basis underlying this trade-off is mainly unknown. Recently, evaluation of 407 winter wheat cultivars for grain and spike characteristics identified geographic as well as temporal trends showing a continuous increase in GPS due to an increase in spikelet fertility. The increase in GN associated with spikelet fertility was negatively associated with GW and the trade-off was largely controlled by attached loci on chromosome 2A (Würschum et al. 2018). In light of these report, and our findings that *GNI-A1* mediates trade-off between GN and GW, we suggest that this continuous temporal trend is controlled to some extent by *GNI-A1.* The increasing frequencies of the impaired *GNI-A1* allele over time has driven an increase in GN through the relief of floret abortion and contributed to the genetic barriers limiting an increase in GW.

GW is proposed to be the outcome of an interaction between the potential GW (sink) (i.e., when the assimilate supply is not limiting grain growth), and the actual supply of assimilate during the grain filling period (source) (Fischer 2011). Removal of distal florets in LDN significantly increased the weight of proximal grains (Fig. 4b). Moreover, removal of spikelets increased weight of G3 (Fig. 4a, Supplementary Fig. S5), indicating that in the durum wheat *cv*. LDN, distal grains are also source-limited. Lower GW in LDN was spatially related to higher fertility (Fig. 3, Supplementary Fig. S2), suggesting that in LDN, assimilate supply is shifted towards distal grains, thereby restricting filling of proximal grains.

Irrespective of source limitation, the DIC allele increased the proportion of proximal “large grains” (Supplementary Fig. S4). Several studies suggested that the negative correlation between GN and GW derives from the large proportion of “small grains” at distal positions, and is independent of any competitive relationship among developing grains (Acreche and Slafer 2006; Ferrante et al. 2017; Ferrante et al. 2015). This proposal is supported by low correlations found between final GW and starch-synthesizing enzymes (Fahy et al. 2018), which strengthens the idea that final GW is determined by developmental processes prior to grain filling (Calderini and Reynolds 2000a; Calderini et al. 2001; Simmonds et al. 2016). Yet, in the current study, carpel weight prior to anthesis was similar, indicating that increased weight of G1 and G2 associated with the DIC allele is due to reduced competition for assimilates supply during the grain filling period.

Haplotype analysis demonstrated the loss of genetic diversity during initial wheat domestication. Subsequently, *GNI-A1* accumulated non-synonymous mutations (G182R, N166D, N105Y) during the evolution of domesticated tetraploid wheat (Fig. 5), suggesting that deliberate human selection targeted GNI-A1. The reduced transcriptional activity found in domesticated emmer wheat (Fig. 6d) indicates that transcriptional activity of GNI-A1 was compromised by the introduction of the G182R substitution in the CTR. A previous study in barley has shown that the transcriptional activating domain of VRS1 is localized to the CTR, with no contribution from the NTR and HD-Zip domains (Sakuma et al. 2013). Therefore, we propose early selection of domesticated wheat varieties with reduced GNI-A1 activity, a trend that was strengthened with the introduction of 105Y to durum varieties released more recently.

Allelic variation of *GNI-A1* showed that all wild emmer accessions carry the functional 105N allele, indicating that the impaired allele is associated with a fitness cost. Wild emmer has relatively heavy, arrow-shaped spikelets (dispersal unit), which fall on the ground after spike disarticulation, and penetrate the dry soil (Horovitz 1998). Thus, with such limited seed dispersal capacities, lower fecundity would reduce intra-species competition and thereby increase fitness. The observed trade-off between GN and GW may suggest that suppression of floret fertility is endorsed in wild emmer due to its positive effect on grain size. Large seeds promote seedling survival and growth (Venable 1992) due to their positive effect on the seedling vigor (Jakobsson and Eriksson 2000). Seedling vigor is a key component in plant ability to compete with neighboring plants (Weiner 1990) and, therefore, promotion of grain size by the suppression of floret fertility is likely to aid establishment of wild emmer seedlings in their natural environment. Hap6 represents a rare allele of *GNI-A1* located in a narrow geographic region around the Sea of Galilee in Northern Israel. Accessions from this haplotype are from the *Judaicum* subpopulation, characterized by tall culms, wide leaves, wide spikes and large grains (Avni et al. 2018; Poyarkova et al. 1991). Hap6 holds unique features such as two copies of *GNI-A1* and asparagine (N) in position 180 of the protein. Copy number variations are associated with changes in gene expression (Stranger et al. 2007) and play a key role in adaptation for a broad range of environments (Díaz et al. 2012; Würschum et al. 2015; Zhu et al. 2014). The two copies of *GNI-A1* identified in Hap6 is a phenomenon evident in a definite geographical area, characterized by wide inter-annual fluctuations in water availability (e.g., 173-859 mm year^−1^; Peleg et al. 2008). Such stochastic environmental fluctuations may act as a driving force for evolution of local ecological adaptations associated with large grains (Metz et al. 2010). The higher GW associated with the Zavitan allele was not associated with reduction in GN (Fig. 6b, c), suggesting that Hap6 introduces functional diversity in *GNI-A1* that increases GW without reducing GN. However, the mechanism by which CNV and/or amino acid substitution promotes GW is yet to be discovered.

In two-rowed barley, substitution S184G in a parallel domain of VRS1 resulted in extreme suppression of lateral florets and promoted GW, by reducing sink competition from non-reproductive organs (Sakuma et al. 2017). Such a prominent phenotypical difference was not observed between the two wild emmer accessions carrying either 180S (TTD140) or 180N (Zavitan) (Fig. S7). Prediction of phosphorylation potential in VRS1 of two-rowed barley indicated hypophosphorylation of VRS1 due to the S184G amino acid substitution in the serine-rich motif of the CTR, which was suggested to prolong protein function throughout plant development (Sakuma et al. 2017). Here, we predicted hypophosphorylation of Hap6 proteins caused by the S180N substitution (Supplementary Fig. S8) and found that it does not compromise transcriptional activity of GNI-A1 in a heterologous yeast system. Considering that the assay reports only transcriptional changes that emanate from structural modifications of the protein, this finding suggests that the S180N substitution may affect GNI-A1 function through interference with regulatory processes. Moreover, the G182R amino acid substitution introduced during wheat domestication, reduced transcriptional activity and may be regarded as fundamental for the interaction of *GNI-A1* with the transcription machinery.

Overall, our findings expand the knowledge of the genetic basis underlying trade-off between GN and GW. Exploration of the tetraploid gene pool suggests that wild emmer possesses a unique *GNI-A1* allele that may facilitate breeding for grain weight without a significant compromise on grain number.

## ACKNOWLEDGEMENTS

We highly appreciate the excellent technical support of the Peleg lab members. Specially, we thank N. Teboul for drawing in Figure 4. This research was supported by the Chief Scientist of the Israel Ministry of Agriculture and Rural Development (grant #12-01-0005), and the U.S. Agency for International Development Middle East Research and Cooperation (grant # M34-037).

