## Supplementary material for "*GNI-A1* mediates trade-off between grain number and grain weight in tetraploid wheat": SI data

Supplementary Tables and Figures

**Supplementary Table S1.** Primers used in this study

| | | | |
|---|---|---|---|
| **QTL mapping** | | | |
| **Locus** | **Position (cM)** | **Forward** | **Reverse** |
| *wmc407* | 0 | GGTAATTCTAGGCTGACATATGCTC | CATATTTCCAAATCCCCAACTC |
| *wmc667* | 10.6 | GAGGAGAGGAAAAGGCAGGCTA | AACTCTTGCGTGTCTCAAACCG |
| *wmc177* | 21.5 | AGGGCTCTCTTTAATTCTTGCT | GGTCTATCGTAATCCACCTGTA |
| *hbg224* | 23.2 | GTCGTCATGCAAACATCTGATC | AACAATCACGTACTAACTGGCC |
| *Xhuj001* | 28 | AGCGTCTTAGTACCCTGCTTG | CGTGTTGGCCATGCATAAAAC |
| *wmc522* | 36 | AAAAATCTCACGAGTCGGGC | CCCGAGCAGGAGCTACAAAT |
| *Xhuj002* | 41.6 | CTACCAAATGTGATGCCCGG | GGGAAGTCAAGCGTGTCTGA |
| *Xhuj17* | 43.6 | GCTTACGGCGTTGATATACACA | GCTGAAAACTGAAACTACTCGC |
| *Xhuj003* | 43.6 | CCTCCTGACTCCTCCCTAAA | TGGTAAACCAAAGGTGATAACG |
| *gwm448* | 46.3 | AAACCATATTGGGAGGAAAGG | CACATGGCATCACATTTGTG |
| *gwm95* | 46.8 | GATCAAACACACACCCCTCC | AATGCAAAGTGAAAAACCCG |
| *Barc201* | 49.9 | GCGTTACTAGCCCAGGATTTACATT | GCGTCGATGTGAGTAGCGGGAAAC |
| *cfa2263* | 49.9 | GGCCATGTAATTAAGGCACA | CTCCCAGGAGTACAGAAGAGGA |
| *hbg340* | 51.9 | AACATGCCGCTATCGGTGTTT | GATAATGCAGTAGCGCCAAAG |
| *gwm473* | 52.5 | TCATACGGGTATGGTTGGA | CACCCCCTTGTTGGTCAC |
| *hbg393* | 53.6 | atctgataacctccttgagcacac | catggactcttcctattctccatc |
| *wmc644* | 53.6 | GACCCTGGTATTCGCACCTCTG | CGTGACGGCCATTACATAGGAG |
| *wmc794* | 55.5 | GTAAACTGGAAAGAAAACGAACCTG | CTATCCACACGTGGAAAAGAAATC |
| *hbg494* | 59.4 | cgtcggtaaatcactactcacg | agctggcgtgttcactaggtat |
| *Xhuj008* | 61.1 | ATGTCCATCAATTGCCTTGG | TGGCATCTCATGGTGACCTA |
| *Xhuj14* | 61.1 | CCCAGAGACTTCCTTGCATT | GGATTGGCGCTTCAAAGTAA |
| *cfa2043* | 64 | CAGCCGAAGAAGGATTTCTG | GAGGCAGGAACTTAGGGGAG |
| *barc5* | 67.1 | GCGCCTGGACCGGTTTTCTATTTT | GCGTTGGGAATTCCTGAACATTTT |
| *hbg268* | 70.8 | agatctgagccgttgattgcat | gtacccattgccatccctacac |
| *gwm312* | 77.9 | ATCGCATGATGCACGTAGAG | ACATGCATGCCTACCTAATGG |
| *gwm294* | 87.2 | GGATTGGAGTTAAGAGAGAACCG | GCAGAGTGATCAATGCCAGA |
| *Xhuj004* | 90.2 | CCGAGTTTACCCCATACCAAT | TGGTAAGATTTTGTTATCGCATTT |
| *wmc181* | 91.3 | TCCTTGACCCCTTGCACTAACT | ATGGTTGGGAGCACTAGCTTGG |
| *Barc122* | 121.2 | CCCGTGTATATCCAGGAGTG | CAGCCCTTGTGATGTGATG |
| *gwm382* | 126.3 | CATGAATGGAGGCACTGAAACA | CCTTCCGGTCGACGCAAC |
| **Fine mapping** | | | |
| **Locus** | | **Forward** | **Reverse** |
| *Xhuj005* | | AAGAAGCTTGTCTCAGTTCCTG | GCGGCAAGGCGAAGGTAA |
| *Xhuj006* | | AGGGTACTACTGCTGGCTCT | CCACAGCGTCACAATCAGAT |
| *Xhuj007* | | GCTTACGGGAGGGAGTATGAG | AATCATACGACAACACCCGATG |
| *Xhuj008* | | ATGTCCATCAATTGCCTTGG | TGGCATCTCATGGTGACCTA |
| *Xhuj009* | | AAGGAGCTGACGTACGTGGT | TGGTAGAGCCCGAATCTGTC |
| *Xhuj010* | | CCCGATGGAGATGAATGAAC | GGGCAGATCCACATATCACC |
| *Xhuj011* | | TCAGCCTCATGCTCAACATC | GTCGGTCATCTCCACAGGTT |
| *Xhuj012* | | GGAGCCCTCCATTCGTCAAA | CGCGCCAGATATAAGCACCA |
| *Xhuj013* | | CAGCAAATGACCCAATCAGA | GGTCCAAGACTGACTGTTGCT |
| *Xhuj014* | | CCCAGAGACTTCCTTGCATT | GGATTGGCGCTTCAAAGTAA |
| *Xhuj015* | | CACTTCTGCTGGCAAGATCA | CCACATCATCTCCCTTAGGC |
| *Xhuj016* | | GGGAACTTGAGCTAGGGGAG | TGATCTCAAAACACCCACCC |
| **Re-sequencing** | **Primer name** | **Primer sequence** | |
| *GNI-A1* | GNI-A1_125 | GTTCCTCACCTCACCACACC | |
| *GNI-A1* | GNI-A1_1263 | ATCGCCGGAGAAAGGATTTA | |
| **Cloning** | | | |
| *GNI-A1* | GNI-A1_EcoR1 | AGTCGAATTCATGGACAACCAGCAGCTCTT | |
| *GNI-A1* | GNI-A1_Sal1 | AGTCGTCGACTCATGCGAAAAACCACTCG | |
| **Copy number variation** | | | |
| *GNI-A1* | GNI-A1_3819F | CTAACCCAAGACCGAATGAGT | |
| *GNI-A1* | GNI-A1_3957R | GTCCCTGTTGCCACAAATCC | |
| TraesCS2A02G134000 | Mg5627_F | GCTTACGGCGTTGATATACACA | |
| TraesCS2A02G134000 | Mg5725_R | GCTGAAAACTGAAACTACTCGC | |

**Supplementary Table S2**. List of accessions used for re-sequencing *GNI-A1* and their corresponding haplotypes

| Accession | Specie | Haplotype |
|---|---|---|
| PI 428069 | *T. turgidum* ssp. *dicoccoides* | Hap05 |
| 13-B-53 | *T. turgidum* ssp. *dicoccoides* | Hap06 |
| Alm-1 | *T. turgidum* ssp. *dicoccoides* | Hap06 |
| ISR-A | *T. turgidum* ssp. *dicoccoides* | Hap06 |
| Zavitan | *T. turgidum* ssp. *dicoccoides* | Hap06 |
| 35-9 | *T. turgidum* ssp. *dicoccoides* | Hap02 |
| PI 428054 | *T. turgidum* ssp. *dicoccoides* | Hap02 |
| PI428054 | *T. turgidum* ssp. *dicoccoides* | Hap02 |
| PI 428077 | *T. turgidum* ssp. *dicoccoides* | Hap02 |
| PI428084 | *T. turgidum* ssp. *dicoccoides* | Hap02 |
| PI538666 | *T. turgidum* ssp. *dicoccoides* | Hap02 |
| Rei-1 | *T. turgidum* ssp. *dicoccoides* | Hap02 |
| HR1-24 | *T. turgidum* ssp. *dicoccoides* | Hap11 |
| PI352322 | *T. turgidum* ssp. *dicoccoides* | Hap11 |
| PI503310 | *T. turgidum* ssp. *dicoccoides* | Hap11 |
| 28-6 | *T. turgidum* ssp. *dicoccoides* | Hap01 |
| 10-209 | *T. turgidum* ssp. *dicoccoides* | Hap01 |
| 12-3 | *T. turgidum* ssp. *dicoccoides* | Hap01 |
| 18-60 | *T. turgidum* ssp. *dicoccoides* | Hap01 |
| 24-39 | *T. turgidum* ssp. *dicoccoides* | Hap01 |
| Citr 17676 | *T. turgidum* ssp. *dicoccoides* | Hap01 |
| MM5/4 | *T. turgidum* ssp. *dicoccoides* | Hap01 |
| PI 428014 | *T. turgidum* ssp. *dicoccoides* | Hap01 |
| PI 428018 | *T. turgidum* ssp. *dicoccoides* | Hap01 |
| PI 428092 | *T. turgidum* ssp. *dicoccoides* | Hap01 |
| PI 481521 | *T. turgidum* ssp. *dicoccoides* | Hap01 |
| PI 538626 | *T. turgidum* ssp. *dicoccoides* | Hap01 |
| PI 554583 | *T. turgidum* ssp. *dicoccoides* | Hap01 |
| PI428025 | *T. turgidum* ssp. *dicoccoides* | Hap01 |
| PI428036 | *T. turgidum* ssp. *dicoccoides* | Hap01 |
| PI538642 | *T. turgidum* ssp. *dicoccoides* | Hap01 |
| KH2/3 | *T. turgidum* ssp. *dicoccoides* | Hap10 |
| PI466946 | *T. turgidum* ssp. *dicoccoides* | Hap10 |
| Am-1 | *T. turgidum* ssp. *dicoccoides* | Hap04 |
| TTD140 | *T. turgidum* ssp. *dicoccoides* | Hap04 |
| PI487264 | *T. turgidum* ssp. *dicoccoides* | Hap03 |
| 36-24 | *T. turgidum* ssp. *dicoccoides* | Hap03 |
| Citr 17675 | *T. turgidum* ssp. *dicoccoides* | Hap03 |
| IG 116184 | *T. turgidum* ssp. *dicoccoides* | Hap03 |
| IG 45494 | *T. turgidum* ssp. *dicoccoides* | Hap03 |
| IG 46323 | *T. turgidum* ssp. *dicoccoides* | Hap03 |
| IG 46476 | *T. turgidum* ssp. *dicoccoides* | Hap03 |
| PI 503316 | *T. turgidum* ssp. *dicoccoides* | Hap03 |
| PI 538700 | *T. turgidum* ssp. *dicoccoides* | Hap03 |
| PI428132 | *T. turgidum* ssp. *dicoccoides* | Hap03 |
| PI466957 | *T. turgidum* ssp. *dicoccoides* | Hap03 |
| 8941 | *T. turgidum* ssp. *dicoccoide* | Hap07 |
| G-929 | *T. turgidum* ssp. *dicoccum* | Hap07 |
| PI 197496 | *T. turgidum* ssp. *dicoccum* | Hap07 |
| PI 298586 | *T. turgidum* ssp. *dicoccum* | Hap07 |
| PI182743 | *T. turgidum* ssp. *dicoccum* | Hap07 |
| PI191091 | *T. turgidum* ssp. *dicoccum* | Hap07 |
| PI264964 | *T. turgidum* ssp. *dicoccum* | Hap07 |
| PI326312 | *T. turgidum* ssp. *dicoccum* | Hap07 |
| PI352352 | *T. turgidum* ssp. *dicoccum* | Hap07 |
| PI352361 | *T. turgidum* ssp. *dicoccum* | Hap07 |
| PI377658 | *T. turgidum* ssp. *dicoccum* | Hap07 |
| PI434995 | *T. turgidum* ssp. *dicoccum* | Hap07 |
| PI532302 | *T. turgidum* ssp. *dicoccum* | Hap07 |
| PI606325 | *T. turgidum* ssp. *dicoccum* | Hap07 |
| PI94637 | *T. turgidum* ssp. *dicoccum* | Hap07 |
| PI94649 | *T. turgidum* ssp. *dicoccum* | Hap07 |
| PI94741 | *T. turgidum* ssp. *dicoccum* | Hap07 |
| TTC 01 79 | *T. turgidum* ssp. *dicoccum* | Hap07 |
| TTC 02 80 | *T. turgidum* ssp. *dicoccum* | Hap07 |
| TTC 03 81 | *T. turgidum* ssp. *dicoccum* | Hap07 |
| TTC 09 88 | *T. turgidum* ssp. *dicoccum* | Hap07 |
| Cappelli | *T. turgidum* ssp.*durum* | Hap08 |
| Gaza | *T. turgidum* ssp.*durum* | Hap08 |
| Pavone | *T. turgidum* ssp.*durum* | Hap08 |
| PI 119327 | *T. turgidum* ssp.*durum* | Hap08 |
| PI 25415 | *T. turgidum* ssp. *dicoccum* | Hap08 |
| PI 626468 | *T. turgidum* ssp. *dicoccum* | Hap08 |
| PI 94633 | *T. turgidum* ssp. *dicoccum* | Hap08 |
| PI254169 | *T. turgidum* ssp. *dicoccum* | Hap08 |
| PI322232 | *T. turgidum* ssp. *dicoccum* | Hap08 |
| PI470739 | *T. turgidum* ssp. *dicoccum* | Hap08 |
| Taganrog | *T. turgidum* ssp.*durum* | Hap08 |
| TTC 10 89 | *T. turgidum* ssp. *dicoccum* | Hap08 |
| PI 225332 | *T. turgidum* ssp. *dicoccum* | Hap08 |
| Abu fashi | *T. turgidum* ssp.*durum* | Hap09 |
| Abu fashi | *T. turgidum* ssp.*durum* | Hap09 |
| Aristan | *T. turgidum* ssp.*durum* | Hap09 |
| Aziziah | *T. turgidum* ssp.*durum* | Hap09 |
| Baio | *T. turgidum* ssp.*durum* | Hap09 |
| C61 | *T. turgidum* ssp.*durum* | Hap09 |
| Coll. Jordan | *T. turgidum* ssp.*durum* | Hap09 |
| Dganit | *T. turgidum* ssp.*durum* | Hap09 |
| Durum 33 | *T. turgidum* ssp.*durum* | Hap09 |
| Durum 39 | *T. turgidum* ssp.*durum* | Hap09 |
| Durum 74 | *T. turgidum* ssp.*durum* | Hap09 |
| Eliav | *T. turgidum* ssp.*durum* | Hap09 |
| Inbar | *T. turgidum* ssp.*durum* | Hap09 |
| Juljulith | *T. turgidum* ssp.*durum* | Hap09 |
| kofa | *T. turgidum* ssp.*durum* | Hap09 |
| Kyperounda | *T. turgidum* ssp.*durum* | Hap09 |
| LDN | *T. turgidum* ssp.*durum* | Hap09 |
| m2787 | *T. turgidum* ssp.*durum* | Hap09 |
| MG26427 | *T. turgidum* ssp.*durum* | Hap09 |
| Muri S 50 3 | *T. turgidum* ssp.*durum* | Hap09 |
| Noa | *T. turgidum* ssp.*durum* | Hap09 |
| PI 119330 | *T. turgidum* ssp.*durum* | Hap09 |
| PI 173480 | *T. turgidum* ssp.*durum* | Hap09 |
| PI 178227 | *T. turgidum* ssp.*durum* | Hap09 |
| PI 24492 | *T. turgidum* ssp.*durum* | Hap09 |
| PI 592019 | *T. turgidum* ssp.*durum* | Hap09 |
| Simeto | *T. turgidum* ssp.*durum* | Hap09 |
| Simhon | *T. turgidum* ssp.*durum* | Hap09 |
| Svevo | *T. turgidum* ssp.*durum* | Hap09 |
| Villemur | *T. turgidum* ssp.*durum* | Hap09 |

**Supplementary Table S3.** Correlation values among yield components in RISL and backcrossed recombinant lines.

| RISL, 2016 | Spikes per plant | Grain weight per plant (g) | Number of spikelets per spike | Number of grains per spike | Number of grains per spikelet | Grain weight (mg) |
|---|---|---|---|---|---|---|
| Spikes per plant | **1.00** | **0.75** | 0.02 | 0.14 | 0.18 | -0.47 |
| Grain weight per plant (g) | **0.75** | **1.00** | 0.46 | 0.55 | 0.48 | -0.39 |
| Number of spikelets per spike | 0.02 | 0.46 | **1.00** | **0.66** | 0.36 | -0.03 |
| Number of grains per spike | 0.14 | **0.55** | **0.66** | **1.00** | **0.94** | **-0.59** |
| Number of grains per spikelet | 0.18 | 0.48 | 0.36 | **0.94** | **1.00** | **-0.72** |
| Grain weight (mg) | -0.47 | -0.39 | -0.03 | **-0.59** | **-0.72** | **1.00** |

| Backcrossed recombinant lines, 2017 | Spikes per plant | Grain weight per plant (g) | Number of spikelets per spike | Number of grains per spike | Number of grains per spikelet | Grain weight (mg) |
|---|---|---|---|---|---|---|
| Spikes per plant | **1.00** | **0.82** | -0.16 | 0.11 | 0.19 | -0.13 |
| Grain weight per plant (g) | **0.82** | **1.00** | 0.15 | **0.47** | **0.49** | -0.19 |
| Number of spikelets per spike | -0.16 | 0.15 | **1.00** | **0.57** | 0.30 | 0.03 |
| Number of grains per spike | 0.11 | **0.47** | **0.57** | **1.00** | **0.96** | **-0.60** |
| Number of grains per spikelet | 0.19 | **0.49** | 0.30 | **0.96** | **1.00** | **-0.70** |
| Grain weight (mg) | -0.13 | -0.19 | 0.03 | **-0.60** | **-0.70** | **1.00** |

Significant correlations ($P \leq 0.05$) are highlighted in bold

**Supplementary Table S4.** Biometric parameters of QTLs affecting grain number and grain weight on chromosome 2A.

| Trait | Year | LOD[a] | Position(cM) | P.E.V.[b] | d[c] | Increasing allele[d] |
|---|---|---|---|---|---|---|
| GPS | 2017 | 15.34 | 61.06 | 0.538 | -10.7 | LDN |
| Stand.dev. | | 2.635 | 2.175 | 0.06 | 1.052 | |
| GPSt | 2017 | 18.81 | 61.12 | 0.608 | -0.32 | LDN |
| Stand.dev. | | 2.782 | 1.483 | 0.054 | 0.028 | |
| GW | 2017 | 20.24 | 61.49 | 0.632 | 7.269 | DIC |
| Stand.dev. | | 2.56 | 0.521 | 0.046 | 0.562 | |
| GW | 2014 | 13.33 | 60.03 | 0.48 | 5.752 | DIC |
| Stand.dev. | | 2.916 | 2.811 | 0.073 | 0.62 | |

**a** LOD scores that were found to be significant when comparing hypotheses $H_1$ (there is a QTL in the chromosome) and $H_0$ (no effect of the chromosome on the trait), using the 1000 permutation test (Churchill and Doerge, 1994).

**b** Proportion of explained variance of the trait.

**c** The additive effect of an allele calculated as one-half of the mean difference between homozygotes with and without the allele.

**d** Parental allele contributing to higher values.

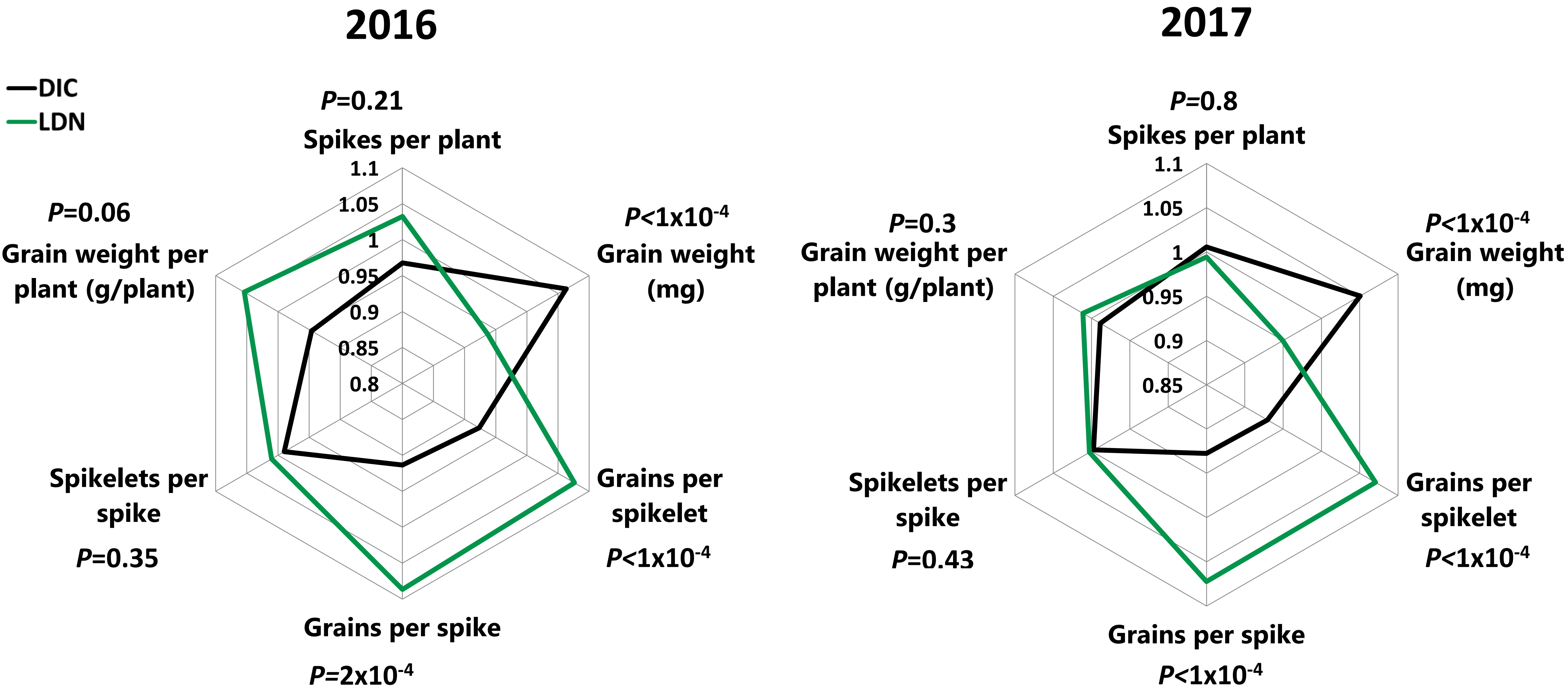

**Fig. S1.** The effect of GW QTL on yield components. **a)** Radar chart demonstrating relative differences in yield components among RISL carrying LDN (*n*=6) or DIC (*n*=7) allele. **b)** Radar chart demonstrating relative differences in yield components among Backcrossed recombinant lines carrying LDN (Green) or DIC (Black) allele (*n*=9). *P* values were determined using student's t test.

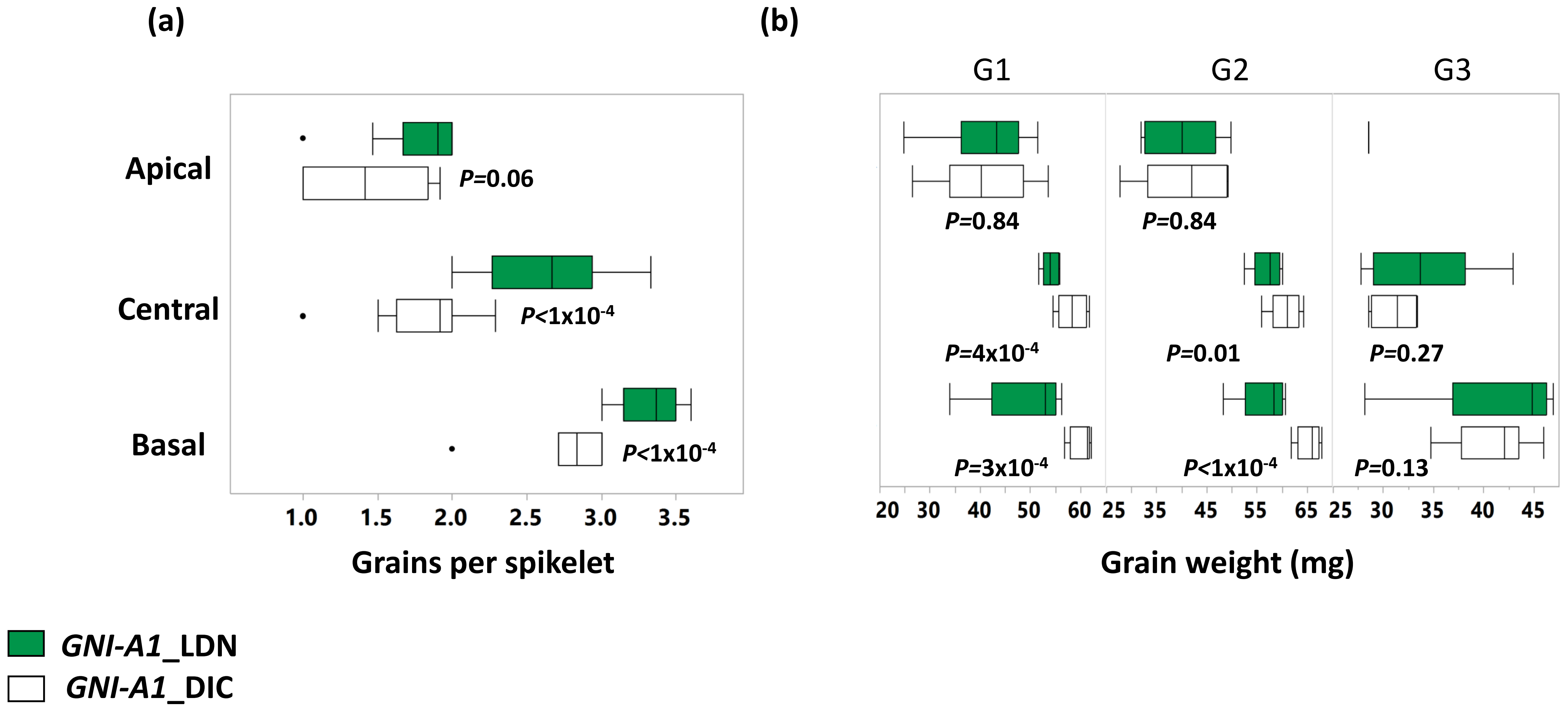

**Fig. S2** Phenotypic characterization of *GNI-A1* locus effect on floret fertility and grain weight. **a**) Number of grains in basal, central and apical spikelets. **b**) weight of particular grains in basal, central and apical spikelets. Green box plot indicates LDN allele. P values were determined using student's t test (*n*=5).

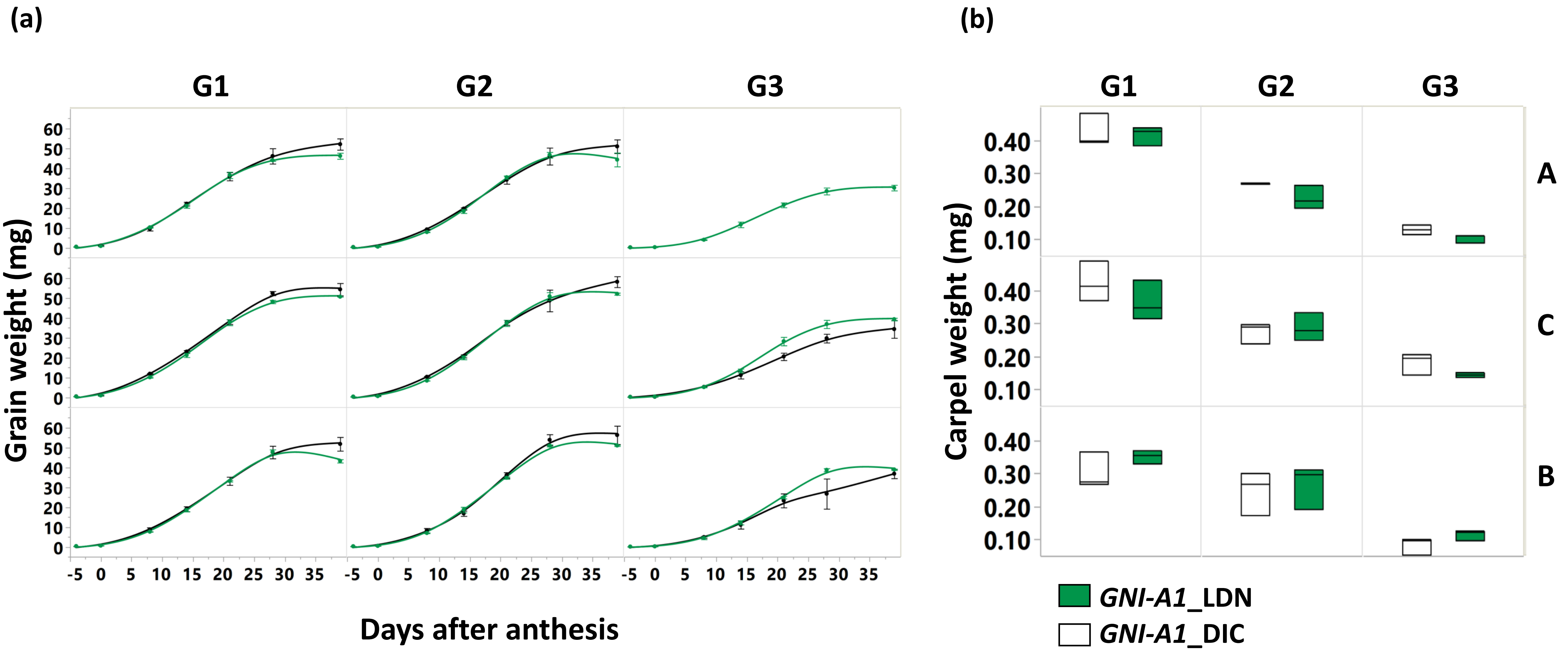

**Fig. S3** Characterization of grain development**. a**) Grain dry weight accumulation in LDN and a near isogenic line (NIL80) from heading to maturity. **b**) Carpel weight at heading. Values are mean of four spikelets per spike section (basal, central, apical) of three spikes (*n*=3).

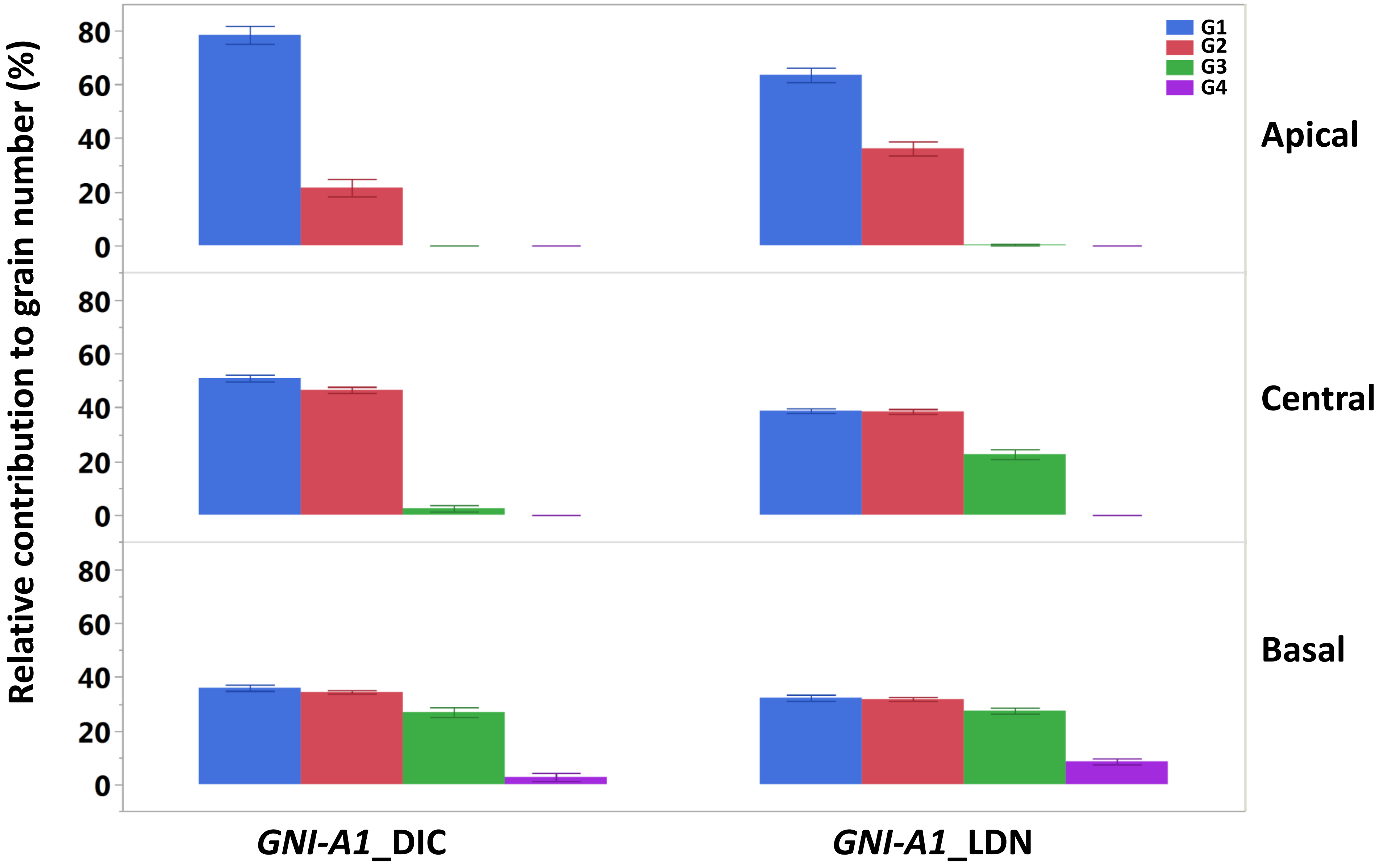

**Fig. S4** Relative contribution of grains from different grain positions to the final grain number. Different colors indicate different grain positions in Basal, Central and Apical spikelets.

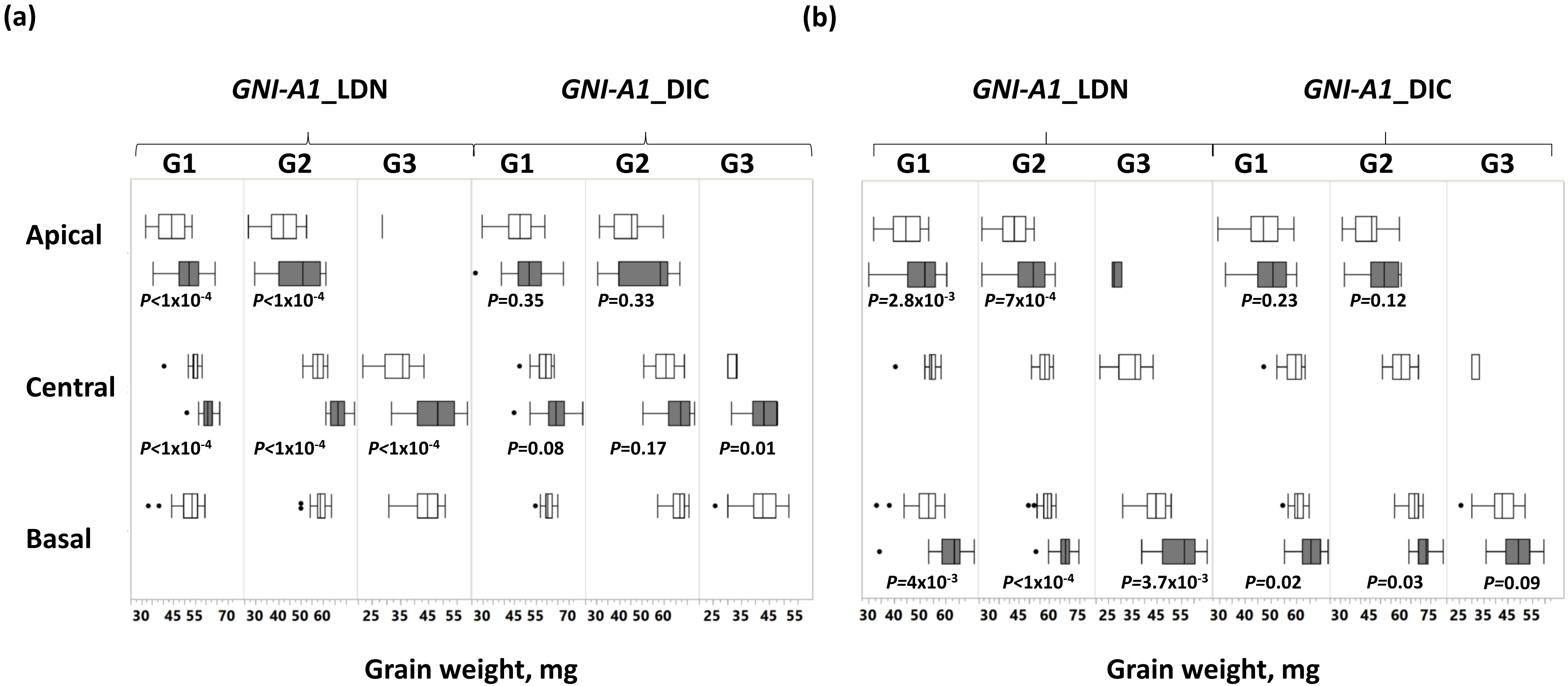

**Fig. S5** Effect of spikelets removal on grain weight in basal, central and apical spikelets of LDN and a Backcrossed recombinant line (63-18). **a)** Weight of particular grains following removal of basal spikelets. **b)** Weight of particular grains following removal of central spikelets. White and gray box plots represent control and treatment (spikelet removal), respectively. P values were determined using student's t test *(n=5)*.

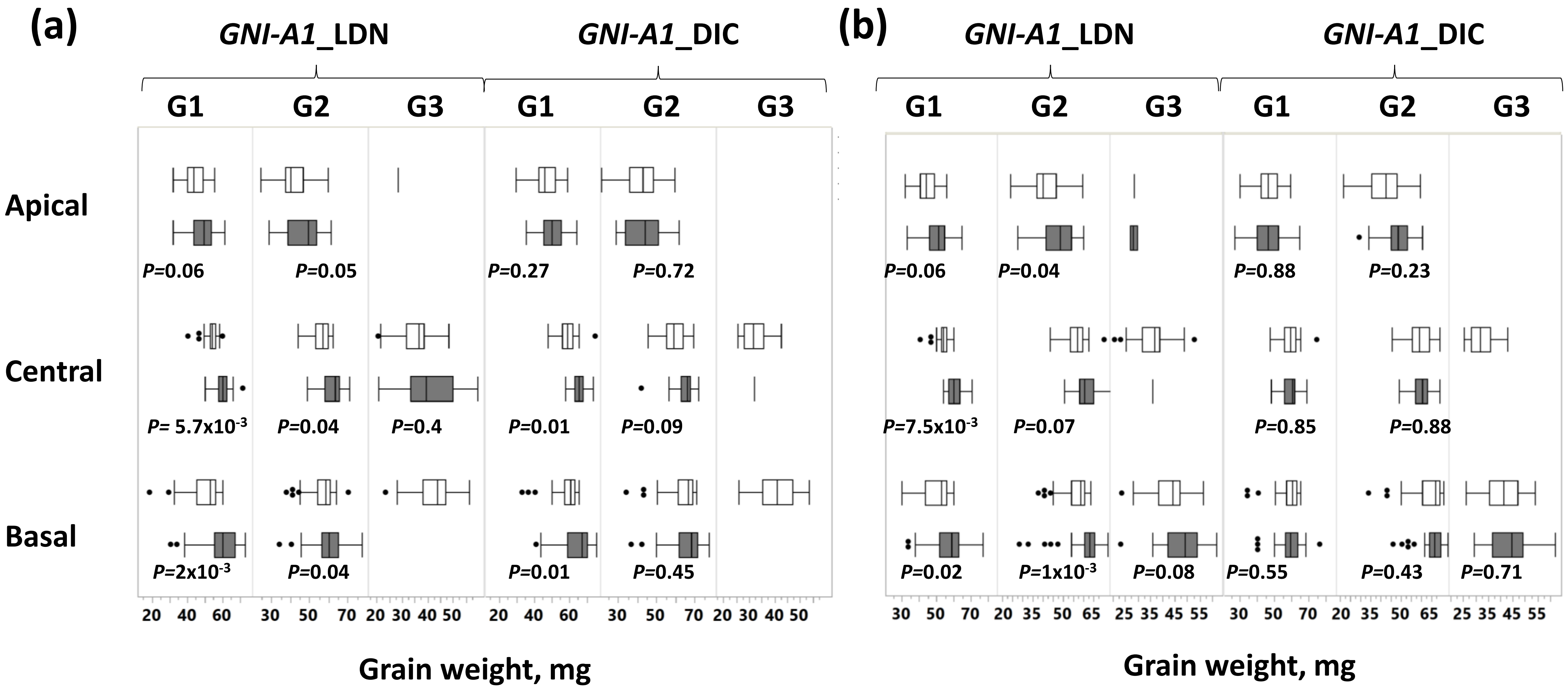

**Fig. S6** Effect of distal floret removal on grain weight in basal, central and apical spikelets of LDN and a Backcrossed recombinant line (63-18). **a)** Weight of particular grains following removal of florets in basal spikelets. **b)** Weight of particular grains following removal of florets in central spikelets. White and gray box plots represent control and treatment (florets removal), respectively. P values were determined using student's t test (*n=5*).

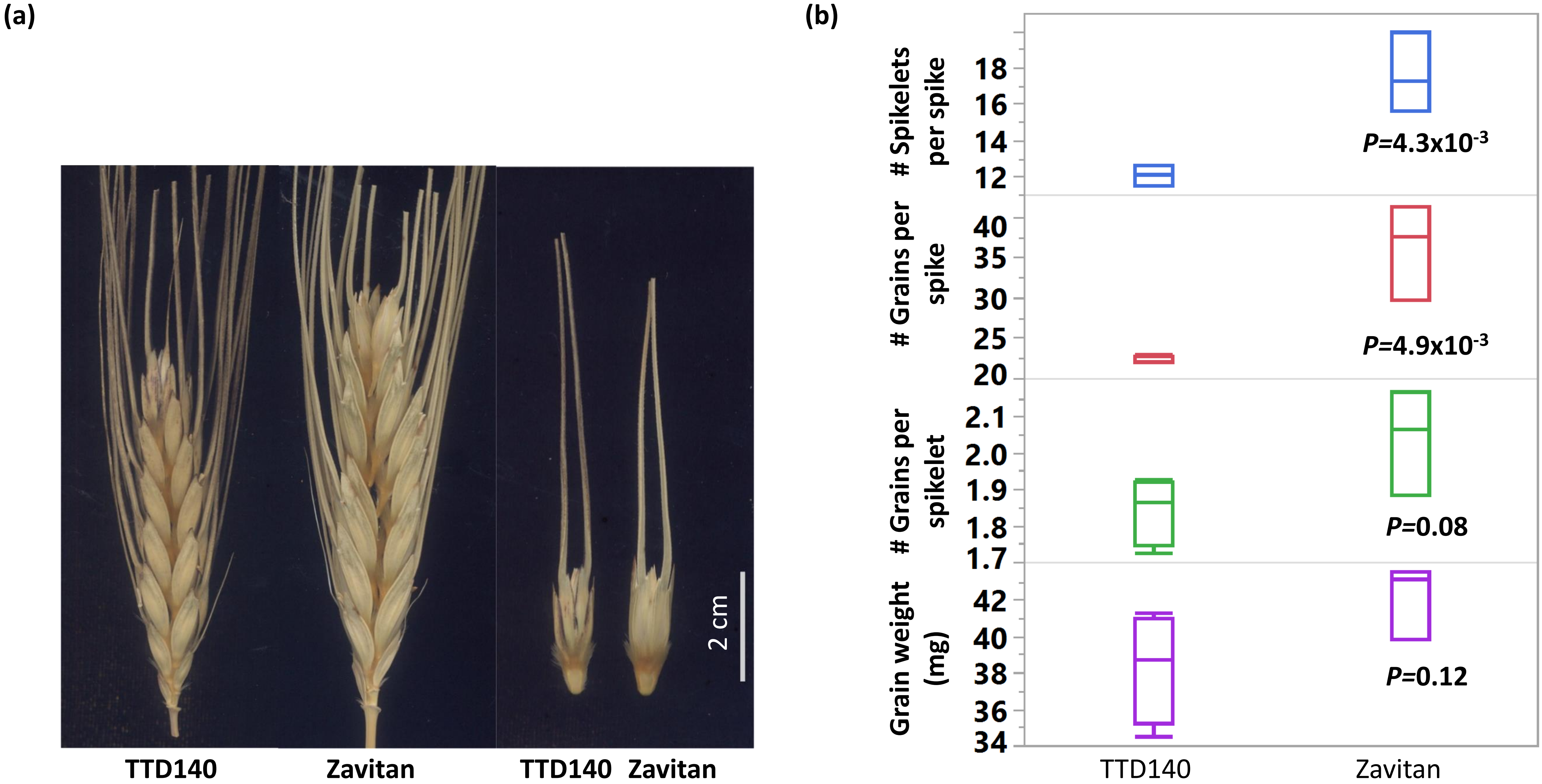

**Fig. S7** Phenotypic characterization of F2 parental lines, TTD140 and Zavitan. **a**) Representative photo of spikes and spikelets of TTD140 and Zavitan. **b**) Characterization of yield components ($n$=4). P values were determined using student's t test.

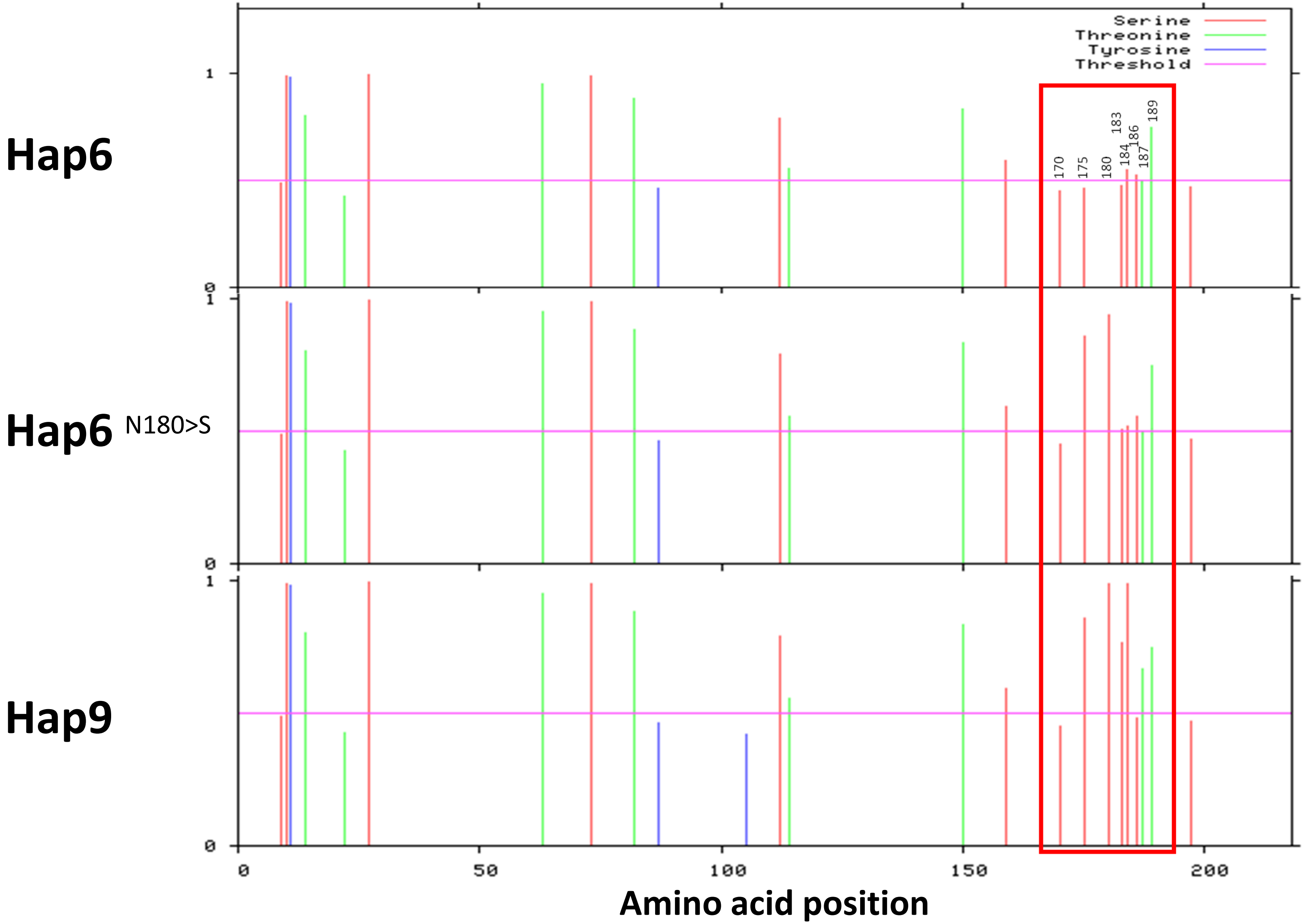

**Fig. S8** Prediction of phosphorylation sites of serine, threonine and tyrosine in GNI-A1 of wild emmer wheat (Hap6), a mutated variant (N180<S) of Hap6 and a durum (Hap9) protein. Red rectangle shows sites where phosphorylation potential was altered. Values above threshold (Blue horizontal line) marks putative sites for phosphorylation.
